# Silver nanoparticle induced immunogenic cell death can improve immunotherapy

**DOI:** 10.1101/2024.08.13.607617

**Authors:** Ara Sargsian, Xanthippi Koutsoumpou, Hermon Girmatsion, Kiana Buttiens, Carla Rios Luci, Stefaan J. Soenen, Bella B. Manshian

**Affiliations:** NanoHealth and Optical Imaging, Department of Imaging and Pathology, KU Leuven, Leuven, Belgium; Translational Cell and Tissue Research Unit, Department of Imaging and Pathology, KU Leuven, Leuven, Belgium; Leuven Cancer Institute, KU Leuven, Leuven, Belgium

## Abstract

Cancer immunotherapy is often hindered by an immunosuppressive tumor microenvironment (TME). Various strategies are being evaluated to shift the TME from an immunologically ‘cold’ to ‘hot’ tumor and hereby improve current immune checkpoint blockades (ICB). One particular hot topic is the use of combination therapies. Here, we set out to screen a variety of metallic nanoparticles and explored their in-vitro toxicity against a series of tumor and non-tumor cell lines. For silver nanoparticles, we also explored the effects of core size and surface chemistry on cytotoxicity. Ag-citrate-5nm nanoparticles were found to induce high cytotoxicity in Renca cells through excessive generation of reactive oxygen species (ROS) and significantly increased cytokine production. The induced toxicity resulted in a shift of the immunogenic cell death (ICD) marker calreticulin to the cell surface in-vitro and in-vivo. Subcutaneous Renca tumors were treated with anti-PD1 or in combination with Ag-citrate-5nm. The combination group resulted in significant reduction in tumor size, increased necrosis, and immune cell infiltration at the tumor site. Inhibition of cytotoxic CD8+ T cells confirmed the involvement of these cells in the observed therapeutic effects. Our results suggest that Ag-citrate-5nm is able to promote immune cell influx and increase tumor responsiveness to ICB therapies.

## 1 Introduction

Tumors characterized by an immunosuppressive tumor microenvironment (TME) are known to exert mechanisms to "hide" from the immune system. TME-derived cytokines, for instance, increase the expression of inhibitory checkpoint molecules (*e.g*., PD-1, CTLA-4) in T cells, which suppresses T-cell antitumor effector functions and, eventually, drives T cells anergic ((1)). To overcome this immunosuppression, one of the most promising therapeutic approaches is immune checkpoint blockade (ICB)s, through so-called immune checkpoint inhibitors (*e.g*. anti-PD1, anti-CTLA-4 antibodies)(2). Many immune checkpoints consist of ligand-receptor interactions, which can be blocked by antibodies or are modulated by recombinant forms of ligands or receptors, preventing checkpoint proteins from binding to their respective partner proteins. In recent years immunotherapy has gained great momentum in cancer treatment. Blocking PD-(L)1 is one of the main areas that has been researched and various antibodies are clinically approved(3). This is mainly because of the expression of PD-(L)1 in peripheral tissues, which can lead to important immune resistance mechanisms within the TME. Therefore, it is presumed that blocking PD-1 signaling affects the effector stage of the immune response. This inhibitory receptor has also been detected on circulating tumor-specific T cells and tumor-infiltrating lymphocytes (TIL). In addition to the high expression of PD-1 on TILs, PD-L1 expression on the tumor cell surface and on regulatory T cells (T-regs) is also present in the TME. Accordingly, blocking PD-L1 will suppress T-reg immunosuppressive activity(4).

In addition to ICB, immunogenic cell death (ICD) is known to play an important role in stimulating the antitumor immune system. ICD occurs when, during apoptotic cell death, cells release damage-associated molecular patterns (DAMPs), which leads to the attraction and activation of immune cells(5). These DAMPs released during ICD include cell translocated calreticulin (CRT), ERp57, adenosine triphosphate (ATP), heat shock protein (HSP) 70/90 and highly mobility group B1 (HMGB1)(6). Several therapeutic strategies, such as some forms of chemotherapy and radiotherapy, have been shown to induce ICD at the tumor site(7).

Nowadays, immunotherapy (IT) is frequently used in clinical settings, however with a response rate of 15 to 20% it is not effective in all patients. Therefore, more and more, combinations of immunotherapy with other therapeutic mechanisms such as chemotherapy are applied in order to achieve higher response rates. In general, these combined strategies are found to improve the survival rate in patients as they increase the immunogenicity of the tumor(8).

Apart from chemotherapy, IT has also been more frequently combined with nanoparticles (NPs) for improved efficacy(9). In particular inorganic NPs have shown potential in eliciting antitumor immune responses, as demonstrated for iron oxide NPs that were found to repolarize tumor-associated macrophages (TAMs)(10), or for doped CuO and ZnO NPs which were found to improve immunotherapeutic regimens(11). While many of these inorganic NPs have been evaluated for their use as contrast agents for biomedical imaging, or as therapeutic agents against infectious diseases(12), the crystal structure of the NPs has been reported to activate NLRP3 inflammasomes(13), and cellular uptake of the NPs causes acid etching of the NP surface, resulting in the release of potentially toxic metal ions and reactive oxygen species (ROS)(14). Several reports have indicated that these properties could be utilized for cancer-selective therapy(15). One type of NP that is commonly researched for biomedical applications, are silver (Ag) NPs. Due to their anti-bacterial properties, Ag NPs are already used in clinical wound dressings or so-called diabetic socks. High levels of Ag NPs will however be harmful to mammalian cells, and the NPs must therefore be used with great care. In view of the latter, it has become highly indispensable to investigate the nano(cyto)toxicity of NPs in general, and Ag NPs in particular(16). One question that remains is to what extent the Ag NPs, already demonstrating preliminary anti-tumor capabilities, possess potent anti-tumor properties and how they may be able to tune the immune system to aid common ICB. It is furthermore important to note that many properties of NPs, such as size, coating, charge and shape can affect their cytotoxicity or therapeutic potency.

The aim of this study was to determine which Ag-NPs formulations, with different coatings and sizes, are able to elicit immune-modulating properties and reveal anti-tumor potency and to what extent they compare to other metallic NPs we have previously tested in-house, including Cu-doped TiO_2_ (17), Fe-doped ZnO(18) and Fe-doped CuO(19). Finally, we aim to evaluate whether Ag NPs can synergistically be combined with ICB for improved therapeutic efficacy in immunologically cold tumors.

## 2 Material and Methods

### 2.1 Nanoparticle characterization

Different coated (polyvinylpyrrolidone (PVP; nanoComposix, USA), citrate (nanoComposix, USA), and polyethylene glycol (PEG; nanoComposix, USA)) and sized (5nm and 50nm) silver (Ag) NPs were used. Dynamic light scattering (DLS) and zeta potential (ZP, mV) of Ag-NPs were measured using the Zetasizer Nano ZS (Malvern, instruments, UK). Ag-NP concentration and size was detected using NanoSight LM10 (Malvern Instruments, Worcestershire, UK). Imaging of Ag-NPs was performed using high resolution transmission electron microscopy (TEM 1400, Jeol, Philips, NL) at magnifications: X8k (500nm Ag-NP), X50k (100nm Ag-NP), X150k (20nm Ag-NP).

### 2.2 Cell culture

Mouse breast adenocarcinoma (4T1), mouse renal adenocarcinoma (Renca), mouse colorectal carcinoma (CT26), human cervical cancer (HeLa), human lung cancer (A549), and mouse mesenchymal stem (mMSC) cell lines were used. All cell lines were obtained commercially (Atcc, Belgium). 4T1, HeLa and A549 cells were cultivated in Dulbecco’s Modified Eagle Medium (DMEM; Gibco, ThermoFisher Scientific, Belgium) supplemented with 10% fetal bovine serum (FBS; Gibco, ThermoFisher Scientific, Belgium) and 1% penicillin/streptomycin (Corning, Belgium). Renca cells were cultivated in Roswell Park Memorial Institute medium (RPMI 1640 – GlutaMax^TM^; Gibco, ThermoFisher Scientific, Belgium) supplemented with 10% FBS, 1mM sodium pyruvate (Gibco, ThermoFisher Scientific, Belgium), 1% non-essential amino acids (HyClone, Fisher Scientific, Belgium) and 1% penicillin/streptomycin (Croning, Belgium). CT26 cells were cultivated in RPMI 1640 supplemented with 10% FBS, and 1%P/S. mMSC were cultivated in DMEM supplemented with 10% FBS and 10% horse serum (HS; Gibco, ThermoFisher Scientific, Belgium). All cell types were incubated at 37°C in a 5% CO_2_ humidified environment.

### 2.3 *In vitro* nanoparticle cytotoxicity evaluation

To analyze nanoparticle cytotoxicity, cells were seeded during their exponential growth phase. A549, Renca, CT26, HeLa, and mMS were seeded at a density of 2000 cells/well, 4T1cells were seeded at a density of 1500 cells/well in sterile 96-well plates (Costar, Belgium) and incubated for 24h at 37°C and at 5% CO2. The next day, cells were treated with increasing concentrations (0, 1 5, 10, 15, 20, 25 μg/ml) of inorganic NPs. Different NP coatings were assessed: polyvinylpyrrolidone (PVP; nanoComposix, USA), citrate (nanoComposix, USA), and polyethylene glycol (PEG; nanoComposix, USA). 4%Fe-Cu, 2%Fe-ZnO, 33%Cu-TiO2 were a part of a collaboration. Serial dilutions of NPs in total cell culture medium (according to the cell type) were prepared per type of differently coated Ag NPs and administered in triplicates. After 24 hours of incubation, the cells were washed twice with Phosphate Buffered Saline (PBS; Gibco, ThermoFisher Scientific, Belgium), followed by staining for further microscopic analysis.

### 2.4 Analysis of viability and mitochondrial oxidative stress

Following exposure to the NPs, cells were washed twice with PBS and incubated for 30 minutes with 100nM Image-IT DEAD green viability stain (ThermoFisher Scientific, Belgium) and 200nM Mito Tracker RED CMXRos (ThermoFisher Scientific, Belgium) in total cell culture medium (100 μL/well). Cells were then washed with PBS and fixed in 2% paraformaldehyde (PFA) (pH = 7,4) for 10 minutes at room temperature (RT). After fixation, the cells were counterstained with 0,2% Hoechst nuclear stain (Hoechst 33342; ThermoFisher Scientific,Belgium) in PBS for 10 minutes. Viability, mitochondrial stress and the generation of reactive oxygen species (ROS) were measured using an automated epifluorescence microscope (INCell Analyzer 2000, GE Healthcare Life Science; Light Microscopy and Imaging Network (LiMoNe), VIB-KULeuven). The microscope was set to detect FITC (viability), dsRed (mitochondrial stress and oxidative stress) and DAPI (cell nucleus). Twelve random image fields were obtained per well with a 20X objective (NA 0.45). Analysis of fluorescent images was performed using the INCell Investigator software (GE Healthcare Life Science). Cell nuclei (DAPI) were first identified and segmented, after which FITC intensity signals and mitochondrial intensity signals were determined and linked, via segmentation, to the cell nucleus of the cell to which they belonged, after which the intensity and the area were determined per cell. Cell viability was then determined as: the number of cells with FITC signal at the cell nucleus; mitochondrial ROS was determined as: the intensity of the mitochondrial network of the cell, and mitochondrial stress as: the total area of the mitochondrial network of the cell. These values were always normalized to the control cells included in each assay to avoid inter-variability between different experiments. Viability, ROS and mitochondrial stress are therefore expressed as percent relative values.

### 2.5 Analysis of cellular morphological changes

After incubation with the different NP formula’s, cells were washed twice with PBS and fixed for 10 minutes with 2% PFA (pH = 7,4) at RT. After fixation, cells were incubated for 1 hour with Actin-stain^TM^ 488 phalloidin (Cytosceleton, Inc, Belgium). After staining, the cells were washed 2x with PBS and counterstained for 10 minutes with 0,2% Hoechst nuclear stain. The actin area was measured using an automated epifluorescence microscope (Operetta CLS, PerkinElmer; Light Microscopy and Imaging Network (LiMoNe), VIB-KULeuven). The microscope was set to detect FITC (actin area), and DAPI (cell nucleus). Twelve Random image fields were obtained per well with a 20X objective (NA 0.45). Analysis was performed using the CellProfiler cell analysis software (Broad Institute, USA). The different cell nuclei were first identified and segmented, after which the FITC signal was determined by means of intensity values. Cell actin area was then determined as the total area of the actin network of the cell, stained with Actin-stain TM 488-Phalloidin. These values were always normalized to the control cells included in each assay to avoid inter-variability between different experiments.

### 2.6 Analysis of immunogenic cell death

Renca cells were seeded in a 12-well plate (Greiner Bio-one, Belgium) with a density of 1,2 x 10^5^ cells/well. Renca cells were incubated with Ag-citrate-5nm NPs at a concentration of 5µg/ml for 24h. Supernatant was discarded and cells were washed 1 time with 1mL PBS. 500µL of 0.25% trypsin (Gibco™ Trypsin-EDTA (0.25%), phenol red, fisher scientific,USA) was added to the cells and incubated for 5 minutes at 37°C. 1mL of cell media was added directly in the wells and mixed, media and cells were then transferred to test tubes. The tubes were centrifuged at 400xg for 5 min and the supernatant was discarded. The cells were washed 1 time with 1x PBS and centrifuged again at 400xg for 5 min, the supernatant discarded, and the cells were stained with recombinant anti-calreticulin antibody (Recombinant Alexa Fluor® 647 Anti-Calreticulin antibody-ER Marker (ab196159), abcam, UK) and MHC class I antibody (MHC Class I (H-2Kd/H-2Dd) Monoclonal Antibody (34-1-2S), FITC, eBioscience™, ThermoFisher science, Belgium) for 1h on ice. Cells were washed with 2mL washing solution (WS; 1xPBS + 1%FBS) and centrifuged at 400xg for 5min. Hereafter, cells were washed with 1xPBS and centrifuged at 400xg for 5min. Cells were re-suspended in 1xPBS and transferred to 1,5mL micro test tubes (total volume not more than 200µL) and detected using image-based flow cytometry (Amnis® ImageStream®X Mk II, Luminex, USA). Analysis was performed using IDEAS application v6.0 (Luminex, USA).

### 2.7 Analysis of cytokine expression

Renca cells were seeded in a 12-well plate (Greiner Bio-one, Belgium) with a density of 1,2 x 10^5^ cells/well. Renca cells were incubated with Ag-citrate-5nm NPs at a concentration of 2µg/ml and 5µg/ml for 24h. Supernatant was discarded and cells were washed 1 time with 1mL PBS. The cells were incubated with brefeldin A (3µg/ml) (Sigma-aldrich, USA)(3µg/mL) for 4h at 37°C. After incubation the cells were washed with 1x PBS and 500µL of 0.25% trypsin (Gibco™ Trypsin-EDTA (0.25%), phenol red, fisher scientific,USA) was added to the cells and incubated for 5 minutes at 37°C. 1mL of cell media was added directly in the wells and mixed, media and cells were then transferred to test tubes. The tubes were centrifuged at 400xg for 5 min and the supernatant was discarded. The cells were washed 1 time with 1x PBS and centrifuged again at 400xg for 5 min, the supernatant discarded. The cells were fixed for 10 minutes with 2% PFA (pH = 7,4) at RT. The cells were washed 1 time with 1x PBS and centrifuged again at 400xg for 5 min, the supernatant discarded. Permeabilization was done using Trition 100x (final concentration 1x in PBS) for 10 min at RT. The cells were washed again with 1x PBS at 400xg for 5min. The cells were incubated with Fc receptor blocking antibody (CD16/CD32 Rat anti-Mouse, Unlabeled, Clone: 2.4G2, BD, fisher scientific) in washing solution (WS;1xPBS+1%FBS) for 30min on ice. After which 2mL of WS was added and centrifuged at 400xg for 5min. The supernatant was discarded and the cells were incubated with primary antibodies (anti-IFN-alpha/beta R2 PE-conjugated (Mouse IFN-alpha / beta R2 PE-conjugated Antibody (FAB1083P), PE, Biotechne), anti-Interferon gamma violetFluor™ 450 (Anti-Interferon gamma antibody (ab253083), violetFluor™ 450, abcam), anti-IL-6 FITC (IL-6 Monoclonal Antibody (MP5-20F3), FITC, eBioscience™, ThermoFisher), anti-IL-12 p35 (IL-12 p35 Monoclonal Antibody (4D10p35), eFluor™ 660, eBioscience™, ThermoFisher); and TNFα (TNF alpha Monoclonal Antibody (MP6-XT22), PE-Cyanine7, eBioscience™ ,ThermoFisher). for 1h on ice (protected from light). After incubation, 2mL of WS was added and centrifuged at 400xg for 5min. Cells were then washed with 1xPBS and re-suspended with 1xPBS and transferred to 1,5mL micro test tubes (total volume no more than 200µL). Antibody-Labelled cells were detected using image-based flow cytometry (Amnis® ImageStream®X Mk II, Luminex, US). Analysis was performed using IDEAS application v6.0 (Luminex, US).

### 2.8 Animal experiment and ethic

Animals were kept in filter top cages with controlled temperature (21 ± 2 °C), humidity (50 ± 10%) and day-night cycle of 12h/12h. Mice received ad libitum standard pellet diet, and water.

Mice were followed up daily. All animal experiments were approved by the ethical commission of animal experiments (ECD) of KU Leuven (approval number: P218/2018 and P219/2018) and were in accordance with principles and procedures in national and European regulations.

### 2.9 Renca subcutaneous tumor model

Renca cells were transduced with a viral vector containing the firefly luciferase gene. Firefly luciferase uses luciferin as a substrate and emits light at 560nm that was detected via non-invasive bioluminescence optical imaging. In this study, 51 female BALB/c mice were used (5 weeks old) (Charles river, Beerse, Brussel). Mice were injected subcutaneously, at the bottom of the right flank, with 1,0 x 10^6^ RencaLuc+ cells

### 2.10 *In vivo* Bioluminescence Imaging (BLI)

Tumor growth was monitored twice a week, using a non-invasive bioluminescence (BLI) optical imaging system (IVIS spectrum; PerkinElmer). For each session, mice were injected intraperitoneally (IP) with 20mM Luciferin (MedChemExpress, USA). The mice were then anesthetized and positioned in the IVIS Spectrum. BLI images were obtained 10 minutes post-administration of Luciferin (medium binning, f stop = 1, excitation time = 20 sec) under general anesthesia with 2% isoflurane inhalation. Regions of interest (ROI) were indicated covering the bioluminescence signal from the tumor. Images were analyzed with LIVING Image processing software (Perkin Elmer, Waltham, MA).

### 2.11 *In vivo* Elastase – Caspase Fluorescence imaging

At the end of the experiment, mice were anesthetized and injected intravenously with Neutrophil Elastase 680 FAST (Neutrophil Elastase™ 680 FAST (NEV11169), Waltham, MA) and NIR-FLIVO 747 Tracer (NIR-FLIVO 747 Tracer *In vivo* Assay, Immunochemistry Technologies). The mice were then positioned in the IVIS Spectrum. Fluorescence images were obtained 4 hours post-administration of the tracers (excitation/emission Elastase: 675nm/720nm, excitation/emission NIR-FLIVO: 747nm/776nm) under general anesthesia with 2% isoflurane inhalation. Regions of interest (ROI) were indicated covering the fluorescence signal from the tumor. Images were analyzed with LIVING Image processing software (Perkin Elmer, Waltham, MA).

### 2.12 Combination treatment protocol

After 14 days, tumor size was measured in Renca xenograft tumor bearing mice using non-invasive BLI. 51 mice in total were divided into 9 groups: group 1 = control group (PBS, n = 9); group 2 = NP-treatment (n = 8); group 3 = immunotherapy anti-PD1 (IT, n = 7); group 4 = combination therapy where only the IT was boosted (NP+IT(boost), n = 7), group 5 = combination therapy where the NP and the IT were boosted (NP (boost) + IT (boost), n = 5), group 6 = anti-CD8 (n = 4), group 7 = NP + anti-CD8 (n = 4), group 8 = anti-PD1 + anti-CD8 (n = 4), group 9 = NP + anti-PD1 + anti-CD8 (n = 4). Immunotherapy was based on monoclonal antibodies against immune checkpoint PD-1 (CD279, *InVivoMAb* anti-mouse PD-1, Biocell). Anti-CD8 was based on monoclonal antibody against CD8 alpha (*InvivoMAb* anti-mouse CD8α, Biocell). Here, mice from groups 3,4,6,7,8 and 9 (anti-PD1, anti-CD8 or both), received intraperitoneal booster injections on day 18, 22 and 26 of 150 µg/mice. NP treatment was based on peritumoral injection of Ag-citrate 5nm NPs on day 14 with a concentration of 20µg/mouse under general anesthesia with 2% isoflurane inhalation. Mice from group 5 received peritumoral booster injection of Ag-citrate-5nm and intraperitoneal booster injection of anti-PD1 on day 18, 22, and 26. Control animals were injected peritumorally with PBS on day 14. Tumor growth was monitored twice a week. All mice were euthanized after 30 days from tumor cell injection.

Another cohort of Renca xenographt bearing mice were treated with higher concentration of Ag-citrate-5nm (50µg/mouse) and anti-PD1 (200µg/mouse). 32 mice were divided into 4 groups: group A = control (PBS, n = 8), group B = NP-treatment (n = 8), group C = anti-PD1 group (n = 8), and group D = NP+anti-PD1 (n = 8). All the groups received booster injections on day 9, 12, 15, and 18 under general anesthesia with 2% isoflurane inhalation.

### 2.13 Tissue preparation

Vital organs and tumors were fixed in 4% PFA (PFA, clinipath, VWR, Belgium). First, organs/tumors were incubated in 15% sucrose (Sigma-aldrich, USA) (in 1x PBS) overnight at 4°C. The next day they were incubated in 30% sucrose overnight at 4°C, after which they were embedded in embedding molds (Peel-A-Way embedding molds, Sigma-Aldrich, USA) using O.C.T (Tissue-Tek, Sakura Finetek, USA). Sections of 10 µm were processed using a cryostat (CryoStar NX70, ThermoFisher Scientific) and fixed on microscope slides (SuperFrost Plus, VWR, Belgium).

### 2.14 Splenocyte isolation and staining

Fresh spleen was collected in 3mL complete medium (CM; DMEM + 10%FBS + 1% peniciline/streptomycin) in a homogenization tube (gentleMACS^TM^ C Tubes, Miltenyi Biotec). Spleen was dissociated using tissue dissociator gentleMACS Octo (gentleMACS^TM^, Miltenyi Biotec). The cell suspension was transferred to a 70µm Nylon strainer (greiner bio-one, Belgium). The suspension was collected in a 50mL tube, transferred to a 15mL tube and centrifuged at 400xg for 5min. After centrifugation, the supernatant was discarded, and the pellet was re-suspended in 3mL red blood cells lysis buffer (NH_4_Cl). Blood lysis was stopped by adding 10mL of CM and then centrifuged at 400xg for 5min. The cells were washed 2 times with 1x PBS and the pellet was re-suspended with PBS and transferred to test tubes: 1/3 of each control sample was taken, mixed and split in 8 test tubes (single antibody staining for matrix compensation) -the remaining cells (2/3) or each control samples were split in 2 tubes (for cocktail A and B antibodies). Cell samples derived from treated spleens were split in 2 test tubes (for cocktail A and cocktail B). The tubes were centrifuged at 400xg for 5 min and the supernatant was discarded. The cells were incubated with Fc receptor blocking antibody (CD16/CD32 Rat anti-Mouse, Unlabeled, Clone: 2.4G2, BD, fisher scientific) in washing solution (WS;1xPBS+1%FBS) for 30min on ice. After which 2mL of WS was added and centrifuged at 400xg for 5min. The supernatant was discarded and the cells were incubated with primary antibodies (cocktail A: anti-CD3 eFluor 450 (CD3e Monoclonal Antibody (145-2C11), eFluor 450, eBioscience™, ThermoFisher), anti-CD19 PE (CD19 Monoclonal Antibody (MB19-1), PE, eBioscience™, ThermoFisher), anti-F4/80 FITC (F4/80 Monoclonal Antibody (BM8), FITC, eBioscience™, ThermoFisher), anti-CD45 PE-Texas Red (CD45 Monoclonal Antibody (30-F11), PE-Texas Red, ThermoFisher); and cocktail B: anti-CD4 APC (CD4 Monoclonal Antibody (GK1.5), APC, eBioscience™, ThermoFisher), anti-CD8α PE-Texas Red (CD8 alpha Monoclonal Antibody (5H10), PE-Texas Red, ThermoFisher), anti-CD69 PE (CD69 Monoclonal Antibody (H1.2F3), PE, eBioscience™, ThermoFisher), anti-CD38 eFluor450 (CD38 Monoclonal Antibody (90), eFluor 450, eBioscience™, ThermoFisher)) for 1h on ice (protected from light). For the matrix compensation, cells from control group were stained separately with the single antibodies. After incubation, 2mL of WS was added and centrifuged at 400xg for 5min. Cells were then washed with 1xPBS and re-suspended with 1xPBS and transferred to 1,5mL micro test tubes (total volume no more than 200µL). Antibody-Labelled cells were detected using image-based flow cytometry (Amnis® ImageStream®X Mk II, Luminex, US). Analysis was performed using IDEAS application v6.0 (Luminex, US).

### 2.15 Immunohistochemistry

10-µm cryostat sections were prepared for immunohistochemical staining. Antigen retrieval was achieved by incubation with proteinase K for 15min. Sections were washed with 1xPBS for 5min and incubated with blocking solution (1x PBS + 10% normal goat serum (NGS, ThermoFisher scientific) + 1% FBS) for 1h. The sections were washed 2 times with 1x PBS and incubated in avidin blocking solution (0.001% avidin (Sigma-aldrich, USA) in 1xPBS) for 20 minutes. The sections were washed twice with 1xPBS and incubated with biotin blocking solution (0.001% biotin (Sigma-aldrich, USA) in 1xPBS) for 20min. After washing twice with 1xPBS, sections were incubated overnight at 4°C with anti-F4/80 antibody (F4/80 antibody | Cl:A3-1, Bio-Rad, USA). Endogenous peroxidases were quenched by incubating the sections in 3% hydrogen peroxide (Acros Organics, Thermofisher Scientific, Belgium) for 20 min. The sections were then washed twice with 1xPBS. Incubation with Anti-Rat-Biotin (Biotin-SP AffiniPure Goat Anti-Rat IgG (H+L), Jackson Immunoresearch, UK) was performed for 1h at room temperature (RT). After washing twice in 1xPBS, the sections were incubated with streptavidin-HRP (HRP-Conjugated Streptavidin, ThermoFisher) for 30min at RT. After washing with 1xPBS, the sections were incubated with tyramide working solution (Alexa Fluor™ 488 Tyramide SuperBoost™ Kit, goat anti-rabbit IgG, Invitrogen, ThermoFisher scientific) by following the manufacturer instructions. Stop reagent (Alexa Fluor™ 488 Tyramide SuperBoost™ Kit, goat anti-rabbit IgG, Invitrogen, ThermoFisher scientific) was added to each section for 2min after which the sections were washed 3 times with 1x PBS. Sections were incubated overnight with anti-CD8 antibody (Novus Biologicals, UK). The sections were washed twice with 1xPBS and incubated with goat-anti-rabbit-polyHRP (Invitroge, ThermoFisher) for 1h at RT. After incubation, the sections were washed once with 1x PBS and incubated for 1h with tyramide working solution (Alexa Fluor™ 594 Tyramide SuperBoost™ Kit, goat anti-rabbit IgG, Invitrogen, ThermoFisher scientific) by following the manufacturer instructions. Stop reagent (Alexa Fluor™ 594 Tyramide SuperBoost™ Kit, goat anti-rabbit IgG, Invitrogen, ThermoFisher scientific) was added for 2min and the sections were washed 3 times with 1xPBS. Hoechst staining was added for 10min at RT and the sections were washed 2 times with 1xPBS. Mounting was performed using fluoromount aqueous mounting medium (Sigma-aldrich, USA) and the sections were covered using cover glass (Rectangular cover glasses, VWR, Belgium). The images were acquired with Vectra Polaris automated system (Akoya Biosciences, USA) and analyzed using open-source software QuPath.

### 2.16 Histology

*Eosin-Hematoxylin staining* - 10 µm cryostat sections were brought to RT for 30 min in Mili-Q water. Tissue sections were washed for 5 min in 1xPBS and stained with hematoxylin (Sigma-aldrich, USA) for 3 min (protected from light). Hereafter, slices were washed with Mili-Q water for 5 min and washed with 80% EtOH (VWR, Belgium) – 0.15% HCl (Acros Organics, Thermofisher Scientific, Belgium) solution for 1 min. Slides were washed again with Mili-Q water for 1 min whereafter they were put in ammonia water (0.3% v/v) for 30 sec. After washing with Mili-Q water for 5 min, they were washed with 95% ethanol for 1 min and stained with eosin (Sigma-aldrich, USA) for 1 min (protected from light). After staining, dehydration was done by washing with 95% ethanol for 5 min followed by two washings with 100% ethanol (5 min each time). After dehydration, clearing of the slides is performed with 100% xylene (clinipath, VWR, Belgium) (5min each). Mounting was done with DPX (Merck KGaA, Germany) and cover glass (Rectangular cover glasses, VWR, Belgium). Images of tissue sections were acquired with the automated Vectra Polaris system (Akoya Biosciences, USA) and analyzed with the open-source software FIJI.

### 2.17 Statistics

Statistical analysis was performed using GraphPad Prism 8.0 (GraphPad Software, USA). Data were presented as mean ± standard error to the mean (SEM). Statistical comparisons between different groups were analyzed using two-way Anova and one-way Anova with the application of Bonferroni correction. The level of statistical significance was indicated when p < 0.05 (*: p < 0.05; **:p < 0.01; ***:p < 0.001, ****:p<0.0001).

## 3 Results and Discussion

### 3.1 AgNP characterization

The different AgNPs were characterized for their size, surface charge and colloidal stability using a variety of methods (**Figure 1A**). The hydrodynamic size of the NPs was measured by dynamic light scattering (DLS), indicating an overall size of approximately 40nm for the citrate and poly(vinyl pyrrolidone) (PVP)-coated 5nm particles. The 50nm core sized particles revealed hydrodynamic diameters of around 72nm for PVP and poly(ethylene glycol) (PEG)- coated NPs, while the citrate-coated ones had a diameter of 96.58 ± 2.80 nm. The apparent discrepancy between the hydrodynamic size and the reported core size by the company can be largely attributed to the difference measurements where hydrodynamic diameter takes into account the core size along with the coating and all solvent ions that are tightly bound to the NP surface, making it much larger than the actual core sizes. To get an idea on polydispersity, the polydispersity index (PdI) of the NPs is also measures, where values of around 0.2 or lower typically indicate a relatively monodisperse NP suspension, while higher PdI values indicate more heterogeneous NP diameters or possible agglomeration occurring. The PdI value clearly indicate that the citrate-coated NPs were the least stable, and likely resulted in some agglomeration in water. This may be due to the dilution of the citrate-stabilized NP in water, where citrate is more loosely bound and citrate molecules may detach from the surface, resulting in potential NP aggregation. Diluting the NPs in biological fluids or using citrate-containing buffers can typically help to avoid this phenomenon(20).

**Figure 1:**
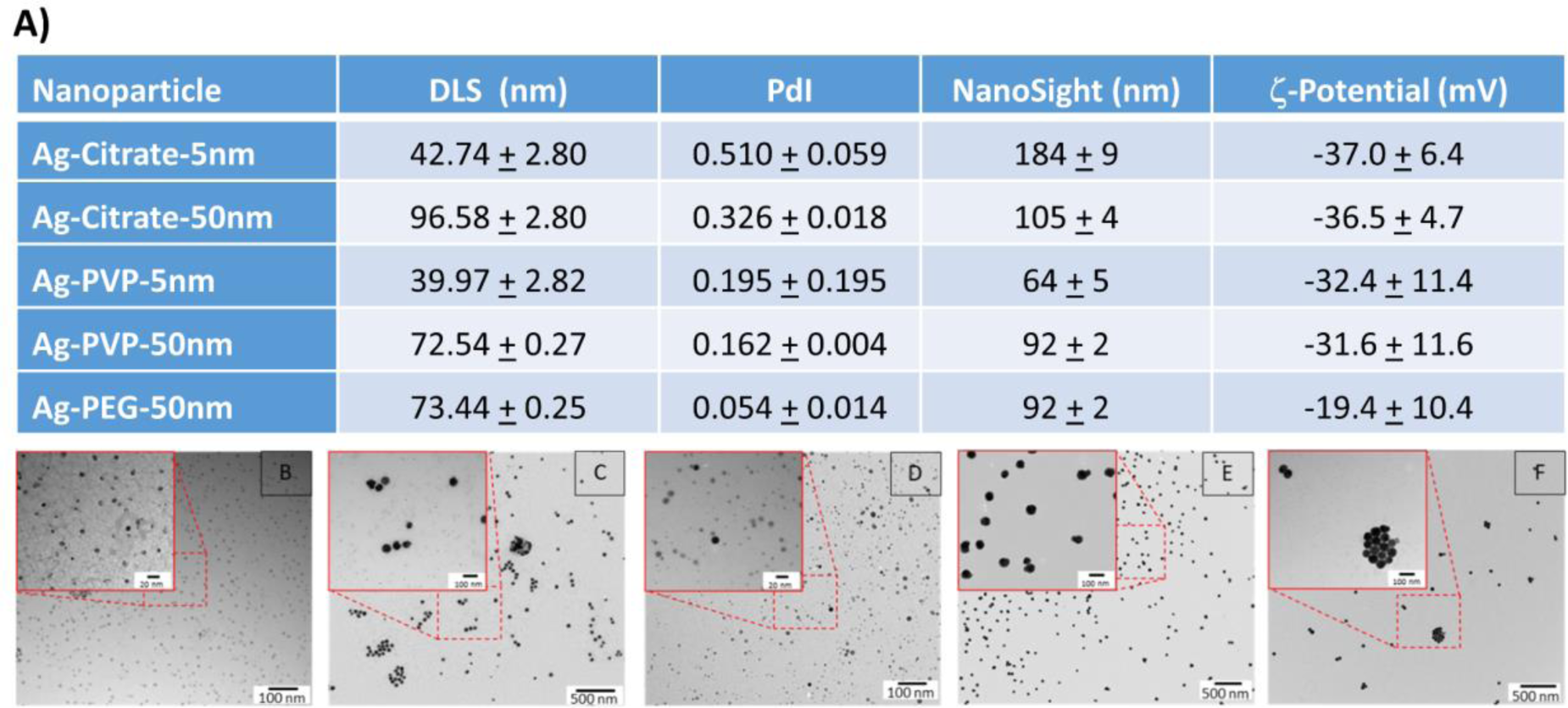
Characterization of silver nanoparticles of different surface coatings and different sizes. A) Table representing the physical characteristics of the different nanoparticles including; hydrodynamic diameter based on dynamic light scattering (DLS) and nanoparticle tracking analysis (NanoSight), polydispersity index (PdI), and surface charge based on the ζ-potential of the different nanoparticles. All data are displayed as mean + standard deviation (n=3). TEM images of B) Ag-citrate-5nm NPs, C) Ag-Citrate-50nm NPs, D) Ag-PVP-5nm NPs, E) Ag-PVP-50nm NPs, and F) Ag-PEG-50nm NPs. The respective scale bars are indicated in both the main image as well as the highlighted (red square) magnified high resolution images.

The poor stability of the citrate-coated NPs was also observed in the nanoparticle tracking analysis (NanoSight LM10), where the sizes of the citrate-coated particles measured exceeded the sizes of their differentially coated counterparts made up of the same core size. The results from the nanoparticle tracking analysis also revealed overall higher sizes than the DLS for each and every particle. This may, in part, be due to the fact that both techniques are not optimally suited to properly analyse very small particles (*i.e.* 5nm core size), where the minimal size for nanoparticle tracking analysis is around 30nm, and is therefore higher than for DLS(21). However, DLS has a much lower peak resolution, and can therefore cannot easily measure heterogeneous samples while nanoparticle tracking analysis can do this more readily(22). The average size measured using both methods may therefore vary more readily.

**Figure 1B-F** displays representative transmission electron microscopy (TEM) images of the different NPs, revealing an overall quite narrow size distribution and core sizes that are in line with those provided by the company.

### 3.2 Ag NPs elicit concentration-dependent ROS and cytotoxicity

Analysis of different cell lines exposed to differently coated and sized Ag-NPs showed varying results in terms of cytotoxicity. The highest effect was seen in Renca cells treated with Ag-citrate-5nm and Ag-PVP-5nm (**Figure 2A**). At the lowest concentration (5µg/mL), both AgNPs variants already induced a highly significant cytotoxic effect (p<0.001). Furthermore, the induced cytotoxicity of Ag-citrate-5nm was accompanied by ROS increases (p<0.01) (**Figure 2B**). Significant changes in mitochondrial area and cell morphology in the Renca cells were also observed for both Ag-citrate-5nm and Ag-PVP-5nm (**Figure 2D**). For the 5nm sized Ag NPs, the mitochondrial and the actin area significantly decreased (p<0.001) in a concentration-dependent manner, starting from a concentration of 15 µg/mL.

**Figure 2.**
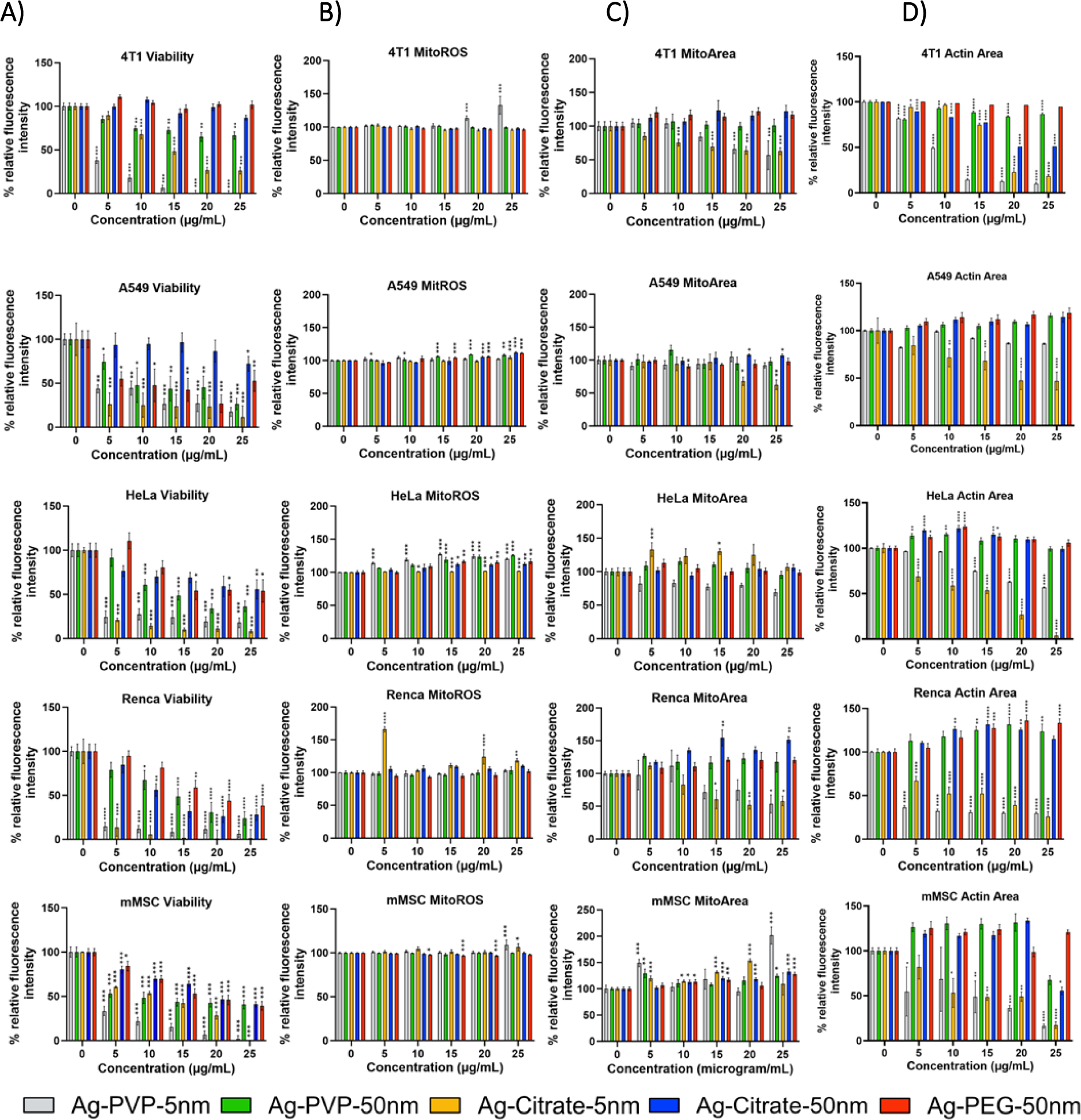
In vitro analysis of Ag NP cytotoxicity in different cells. Representative bar graphs generated for the indicated cells (4T1, A549, Renca, HeLa) and mesenchymal stem cells (MSC). A) Effect on the viability of cells. B) Effect on mitochondrial ROS production. C) Effect on the mitochondrial area. D) Effect on cellular morphology. The data are presented for cells exposed to different inorganic NPs at the indicated concentrations for 24 hours and expressed relative to the level of untreated control cells (100%). The data are gathered from three independent experiments (n=3). The level of significance was indicated when appropriate (*:p< 0.05; **:p< 0.01; ***:p < 0.001; ****:p<0.0001).

In all cases, the smallest Ag-NPs induced the highest cytotoxic effect. As smaller NPs have higher surface area-to-volume ratios, this allows them to release more Ag^+^ ions and therefore the kinetics of Ag^+^ release as well as the extent of surface available for interaction with its physiological environment is higher for smaller NPs(23). Besides their size, their coating also influences Ag^+^ release. Prasad *et al*. previously showed that citrate coated AgNPs resulted in higher levels of intracellular Ag^+^ compared to PVP coated AgNPs(24). In our study, the influence of the coating is less clear, where the toxicity of PVP or citrate-coated 5nm NPs is cell-dependent. This may be attributed to the fact that apart from controlling NP degradation kinetics, the coating will also play a major role in the extent of cellular uptake levels of the NPs. Cellular uptake is in turn highly cell type-dependent as it will depend on the degree of cell cycle progression or total cellular surface area(25). The Renca cells showed the highest sensitivity to Ag-citrate-5nm and Ag-PVP-5nm. At the lowest concentration (5μg/mL), these AgNPs induced a significant decrease in viability, though only the Ag-citrate-5nm induced ROS-mediated cell death. Our data are consistent with those of previous studies done by Gliga *et al*. where it was shown that small AgNPs were cytotoxic to human lung cells and that AgNPs with a size of less than 20nm in diameter induced high levels of ROS production(26). The lack of ROS-mediated cytotoxicity in Renca cell line by Ag-PVP-5nm suggests another mechanism to trigger cell death without increasing mitochondrial ROS production. One possible mechanism is the leakage of cytochrome c from the mitochondria. Ott *et al*. showed that loosely bound cytochrome c can be released from the outer mitochondrial membrane by electrostatic charge (27). Alternatively, cytoplasmic ROS production can induce various cellular damages eventually leading to cell death, while we only took mitochondrial ROS production into account. We therefore suggest further investments are needed to fully understand the mechanism of toxicity in different cell lines induced by differently coated AgNPs. Furthermore, we showed that both AgNPs, Ag-citrate-5nm and Ag-PVP-5nm, significantly reduced the mitochondrial area of Renca cells in a dose dependent manner. This suggests that both AgNPs damage mitochondria, leading to cell death. Changes in cell morphology induced by Ag-citrate-5nm and Ag-PVP-5nm were also dose dependent.

### 3.3 Ag NPs induce higher cytotoxicity than Fe-doped CuO, ZnO, of Cu-doped TiO_2_

Next, we compared the toxicity of the Ag-citrate-5nm to other formulations that had been previously tested in-house and revealed potent anti-tumor immunity (Fe-doped CuO, Fe-doped ZnO and Cu-doped TiO_2_)(17,18). The Ag-citrate-5nm revealed higher levels of toxicity than any of the other formulations in all three cancer cell lines tested (CT26, Renca, 4T1) (**Figure 3**).

**Figure 3.**
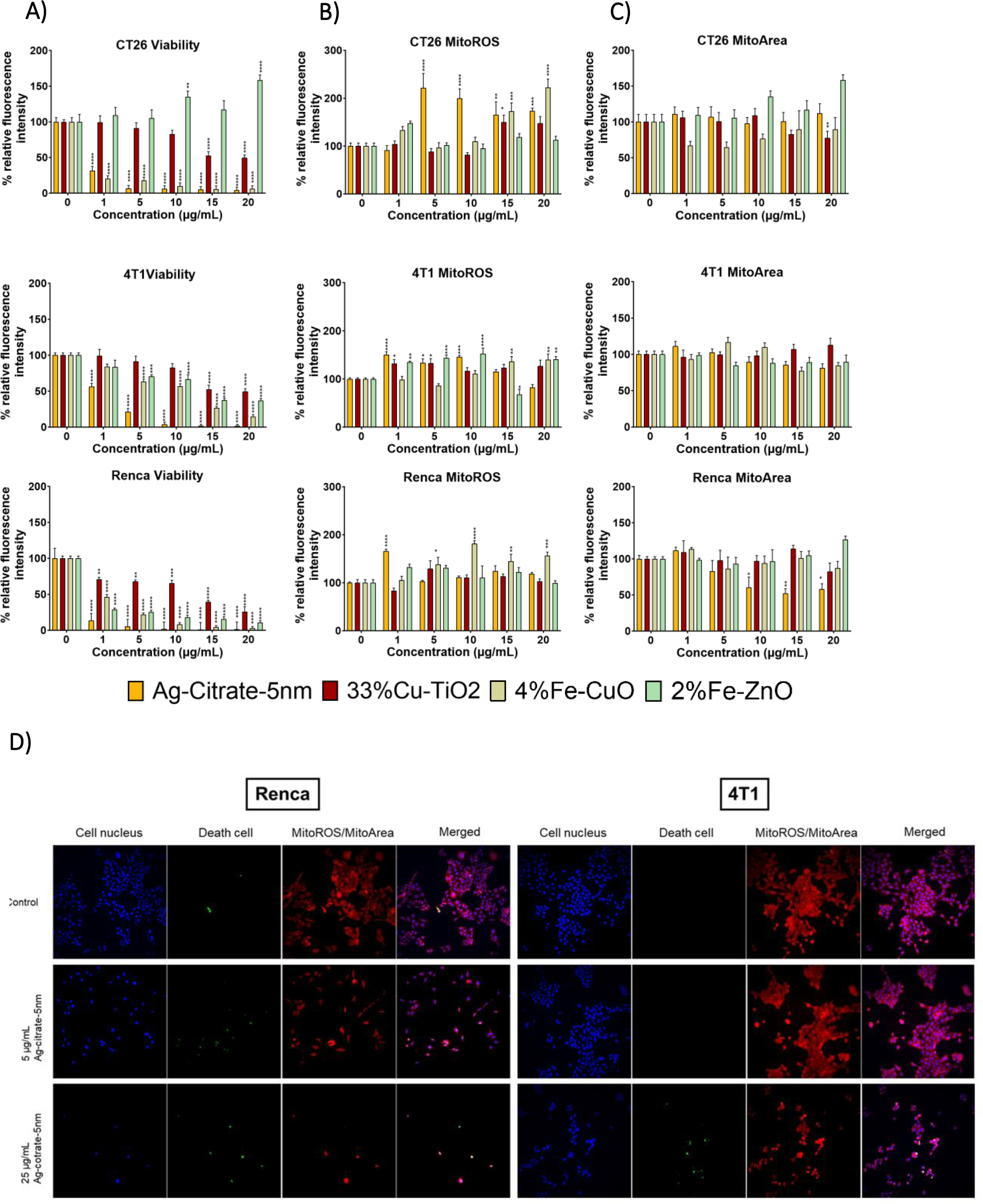
In vitro cytotoxicity of different NP types against murine cancer cells. Representative bar graphs generated for the indicated cells (CT26, 4T1, and Renca. A) Effect on the viability of cells. B) Effect on mitochondrial ROS production. C) Effect on the mitochondrial area. The data are presented for cells exposed to different inorganic NPs at the indicated concentrations for 24 hours and expressed relative to the level of untreated control cells (100%). The data are gathered from three independent experiments (n=3). D) Representative high-content images of Renca (left panel) and 4T1 (right panel) cells stained with the indicated dyes to measure cell death or mitochondrial ROS for control conditions (top row), or cells exposed to Ag-citrate-5nm at 5 µg/ml (middle row) or at 25 µg/ml (bottom row) for 24 hrs. The level of significance was indicated when appropriate (*:p< 0.05; **:p< 0.01; ***:p < 0.001; ****:p<0.0001).

The data reveals that Fe-doped CuO exert the second highest level of cytotoxicity, followed by Fe-doped ZnO and lastly by Cu-doped TiO_2_. The results are somewhat in line with expectations, where overall, it would be expected that pure CuO and ZnO would be more toxic than AgNPs as the former particles possess far higher degradation kinetics. However, the Fe-doped NPs here have been shown to slow down NP dissolution and hereby reduce particle toxicity(28). The particular formulations described here were found to exert the highest level of cancer cell selective toxicity, where they could kill tumor cells under conditions where normal cells remained unaffected(18). While the Ag-citrate-5nm NPs did display higher toxicity in Renca than in MSCs, this formulation has not been optimized for tumor-selective toxicity as it is far more difficult to define conditions through which NP dissolution kinetics can be controlled. For the Cu-doped TiO_2_ NPs, the results revealed hardly any cytotoxicity, which is in line with our previous reports(17). For the Cu-doped TiO_2_ NPs, no cancer cell selective toxicity had been found, but the NPs had been shown to be potent activators of dendritic cells, driven largely by the crystal structure of the largely insoluble TiO_2_ matrix, combined with cancer cell death triggered by released Cu ions. In addition to AgNPs, 4%Fe-CuO NPs induced ROS mediated cell death in Renca cells. Although other inorganic NPs induced cytotoxic effects in Renca, 4T1 and CT26 cell lines, the effect was not ROS mediated.

### 3.4 AgNPs induce immunogenic cell death and promote inflammatory cytokine secretion

As the Ag-citrate-5nm displayed potent toxicity against Renca cells, we examined whether the NP-mediated cell death could evoke immune responses that would promote anti-tumor immunity. For that purpose, initially, we assessed ICD levels from cells exposed to Ag-citrate-5nm and compared them to those from cells exposed to Fe-doped CuO, Fe-doped ZnO and Cu-doped TiO_2_ particles in CT26, 4T1 and Renca cell lines. Our results showed that ICD levels were highly dependent on NP- and cell type. In line with the cytotoxicity results the Ag-citrate-5nm and Fe-doped CuO NPs were the most potent in inducing calreticulin translocation, while both Fe-doped ZnO and Cu-doped TiO_2_ NPs did not cause any significant effects (**Figure 4B**). Next, we explored the effect of varying the Ag NP properties on the induction of ICD in Renca cells. For this end, cells were incubated with the differently sized and coated AgNPs and analysed for the cell surface translocation of calreticulin, a potent marker of immunogenic cell death (ICD). Through ICD, dying cells can release damage associated molecular patterns (DAMPs) to attract and activate dendritic cells to initiate immunogenic cell death(29). It is known that calreticulin, one of the DAMPs and also one of the most abundant proteins in the endoplasmic reticulum (ER), is translocated from the ER to the cell surface due to ROS-mediated ER stress. Under the conditions used, the larger NPs (50nm) did not evoke any ICD, which is in line with the reduced cytotoxicity observed for these larger particles (**Figure 4C**). The biggest effects were clearly observed for the Ag-citrate-5nm, followed by Ag-PVP-5nm. The difference between these two may be related to differences in the level of ROS, where calreticulin translocation and induction if ICD in general have been heavily linked with ROS induction levels(30).

**Figure 4.**
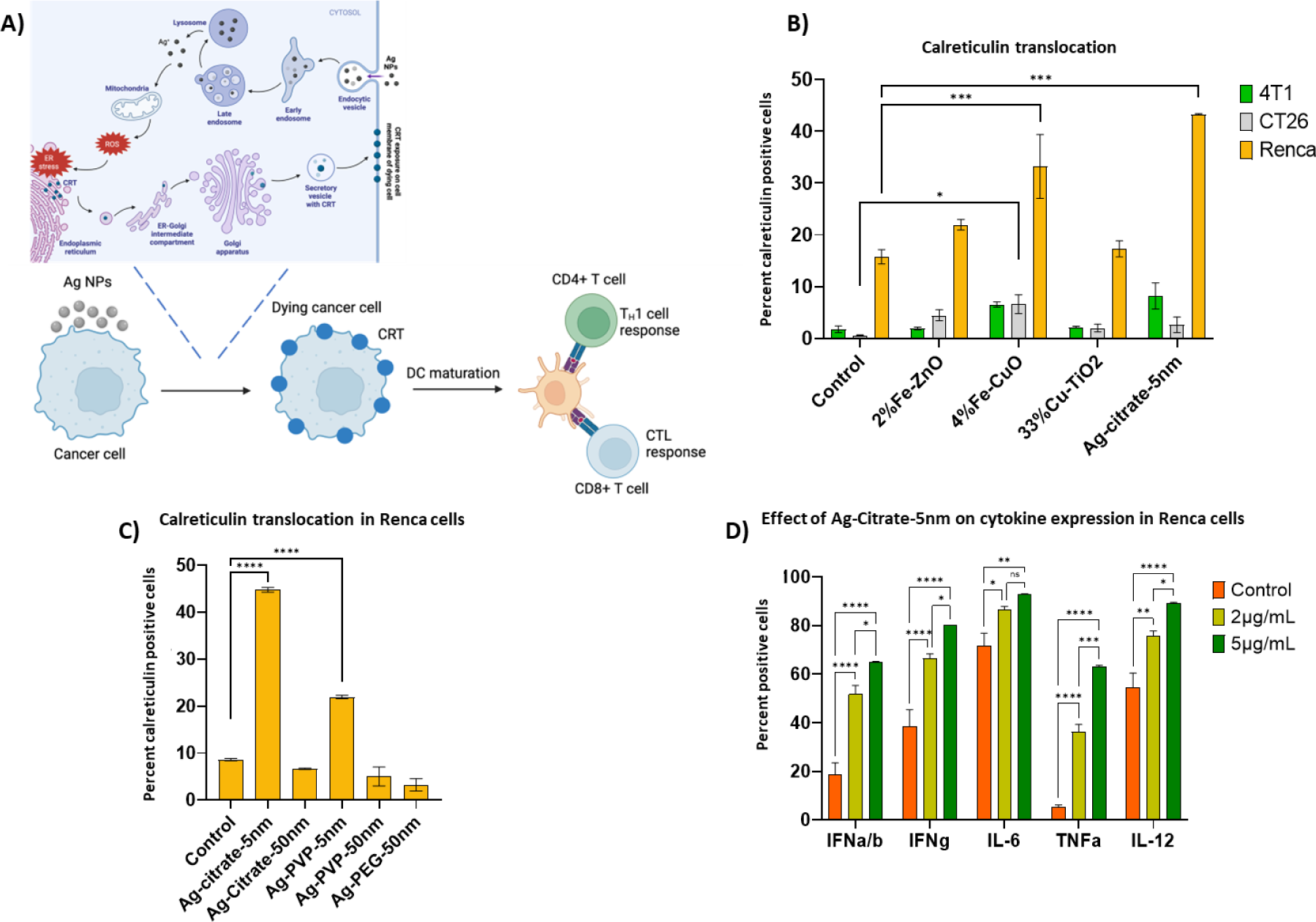
Immunostimulatory effects of Ag NPs evaluated in vitro. A) A schematic describing the mechanism of AgNP induced immunogenic cell death. B) Histograms displaying the level of surface-located calreticulin in Renca, 4T1 and CT26 cells exposed to Ag-citrate-5nm (5µg/ml), 2%Fe-ZnO (15µg/mL), 4%Fe-CuO (5µg/mL), and to 33%Cu-TiO_2_ (30µg/mL) for 24 hours. C) Histograms displaying the level of surface-located calreticulin in Renca cells exposed to differently coated and sized AgNPs at a concentration of 5μg/mL for 24 hrs as measured by ImageStream analysis. D) Cytokine expression in Renca cells after exposure to Ag-citrate-5nm in vitro for 24h. Results are presented as mean + SEM (n=3) in percentages of calreticulin detected on the cell surface or of intracellularly expressed cytokines. The level of significance was indicated when appropriate (*:p< 0.05; **:p< 0.01; ***:p < 0.001; ****:p<0.0001).

It should be noted that in this study, only calreticulin exposure on the cell surface was investigated as a potent ICD marker. In addition to calreticulin, other agents that can be used as ICD markers (*e.g.* ATP, HGMB1, ANXA1, and type 1 IFN) are needed to ensure proper stimulation of anti-tumoral immunity. To better understand the antitumor effect of Ag-citrate-5nm in Renca cells, we therefore examined cytokine expression by exposing the cells to Ag-citrate-5nm at concentrations of 2µg/ml and 5µg/ml. To see the cytokine expression in the cells, we blocked the release 4 hours before staining with Brefeldin A, which inhibits protein transport from the endoplasmic reticulum to the Golgi complex. Our results showed that there was a dose-dependent significant increase in the expression of cytokines such as IL-6, IFNα/β, IFNγ, IL-12 and TNFα (**Figure 4D**).

The cytokines evaluated are all linked to tumor immunity through various mechanisms. Firstly, IL-6 protects cancer cells from therapy-induced DNA damage and oxidative stress by facilitating repair and induction of antioxidant and pro-survival metabolic pathways(31). Since we showed that Ag-citrate-5nm were able to induce cell death mediated by ROS, it further triggers the Renca cells to significantly increase the expression of IL −6 as a defense mechanism to increase antioxidants and induce cell proliferation and reduce cell apoptosis. This is in line with the study of Wu *et al*. which showed that the expression of IL −6 was positively associated with treatment resistance in hormone-resistant prostate cancer(32). Secondly, we also looked at the expression of the IFNα/β receptor (IFNAR2), to which type I IFN can bind(33). The increased signaling through IFNAR2, depends heavily on whether IFNα or IFNβ is binding to the receptor. IFNβ has been shown to be responsible for prosurvival responses(34), while in contrast IFNα induces an antisurvival response in cancer cells by triggering antiproliferative signals and stimulating the cytotoxic activity of various immune cells(35).

A third cytokine is IFNγ, which is mainly produced by natural killer (NK) and T cells. It promotes immune cell activation, maturation, proliferation, cytokine expression and effector function. It also induces antigen presentation, growth arrest, and apoptosis in tumor cells(36). The effect of IFNγ, on immune cells can however be both stimulatory or anti-inflammatory. It can stimulate anti-cancer immunity through CD8^+^ T cell expansion, CD4^+^ T cell polarization into Th1, polarization of myeloid cells toward inflammatory dendritic cells or tumor associated macrophages(37). However, chronic exposure to IFNγ may deplete and impair T cell function by inducing the expression of IDO and PD-L1, directly inhibiting T cell formation, proliferation, clonal diversity, and T cell maintenance. As with immune cells, it can have a dual effect on tumor cells. It exerts an antitumor effect by directly inhibiting tumor growth through apoptosis and necroptosis and growth arrest through senescence while it can increase MHC-I on the tumor cell surface and hereby increase antigenicity(38). However, IFNγ, has also been shown to have tumor-promoting effects in breast cancer and glioblastoma by maintaining survival and stem cell formation, decreasing antigen presentation and enhancing immunosuppressive molecules(39,40).

A fourth cytokine we studied that has pro-inflammatory and anti-tumor properties is IL-12. In the TME, IL-12 is produced and secreted by antigen-presenting cells, which is crucial for the recruitment and effector functions of cytotoxic T and NK cells, while in CD4^+^ T cells it enhances the transcription of T-bet, which increases the expression of Th1-specific chemokines and cytokines to polarize CD4^+^ T cells into activated Th1 cells(41).

The last cytokine we studied was TNFα, which can bind to its 2 receptors (TNFR1 and TNFR2), through which TNFα family members exert their effects. The resulting effects are highly dependent on the concentration of TNFα produced, as well as other possible cytokines in the TME, as TNFα can induce either cell death or alternatively promote cell survival(42). TNFα expression may also serve as a mechanism of immune defense by dedifferentiation of cancer cells and associated loss of immunogenicity(43).

Taken together, the significant induction of various cytokines and receptors by the Ag NPs can trigger a variety of possible responses, which can be either pro- or anti-tumor immunity. However, based on the in vitro data, the use of Ag-citrate-5nm holds a high potential for remodeling the TME and hereby influencing the efficacy of ICB.

### 3.5 Ag NPs synergistically impede tumor growth together with anti-PD1 ICB

Given the potency of the Ag-citrate-5nm NPs on Renca cells *in vitro*, we evaluated the therapeutic efficacy of the particles in a subcutaneous syngeneic luciferase-expression (luc^+^) Renca tumor model. In a first study, animals were treated either with PBS (control), local peritumoral administration of Ag-citrate-5nm, intraperitoneal administration of anti-PD-1 antibody, or a combination of both Ag NP and ICB. Please note that the NPs were injected locally (peritumoral) to obtain high local accumulation. While intravenous administration is typically preferred in clinical settings, this is associated with low accumulation in the tumor and thus low therapeutic efficacy(44). Immunotherapy was always given through 3 repeated doses, each with 4 days in between. In a last combination group, Ag-citrate-5nm was also administered a total of 3 times, with 4 days in between. Tumor growth was then monitored through optical imaging, revealing that single treatments led to higher total tumor flux at day 30 while the combination of NP+anti-PD1(boost) and NP (boost) + anti-PD1 (boost) resulted in a significant reduced lower tumor growth rate (p < 0.0001) (**Figure 5 A**). At day 30, the level of neutrophil elastase and caspase activity at the tumor site were measured through optical imaging. The results showed that the elastase activity in control group (p < 0.01) and the combination group (p < 0.001) was significant higher compared to the anti-PD1 group. Besides, the elastase activity in the combination group was slightly higher than the control group but the difference was not significant (**Figure 5 C**). Tumor cell staining showed that the combination group significantly increased the calreticulin translocation (**Figure 5D**) (p < 0.05). Overall, we could only determine a significant reduction in tumor growth rate in the combination group of Ag-citrate-5nm with anti-PD1 compared to other groups. This effect might be explained by the fact that Ag-citrate-5nm is capable of inducing calreticulin translocation and pro-inflammatory cytokine expression, as shown in our previous *in vitro* experiment, and when combined with anti-PD1, anti-tumoral immune response is triggered, but not strong enough to induce a potent anti-tumor response as a monotherapy. While the tumors did not show a clear response to ICB as a monotherapy, indicating their high level of immunosuppression or lack of tumor infiltrating immune cells, the combination with Ag-citrate-5nm was able to overcome this problem.

**Figure 5.**
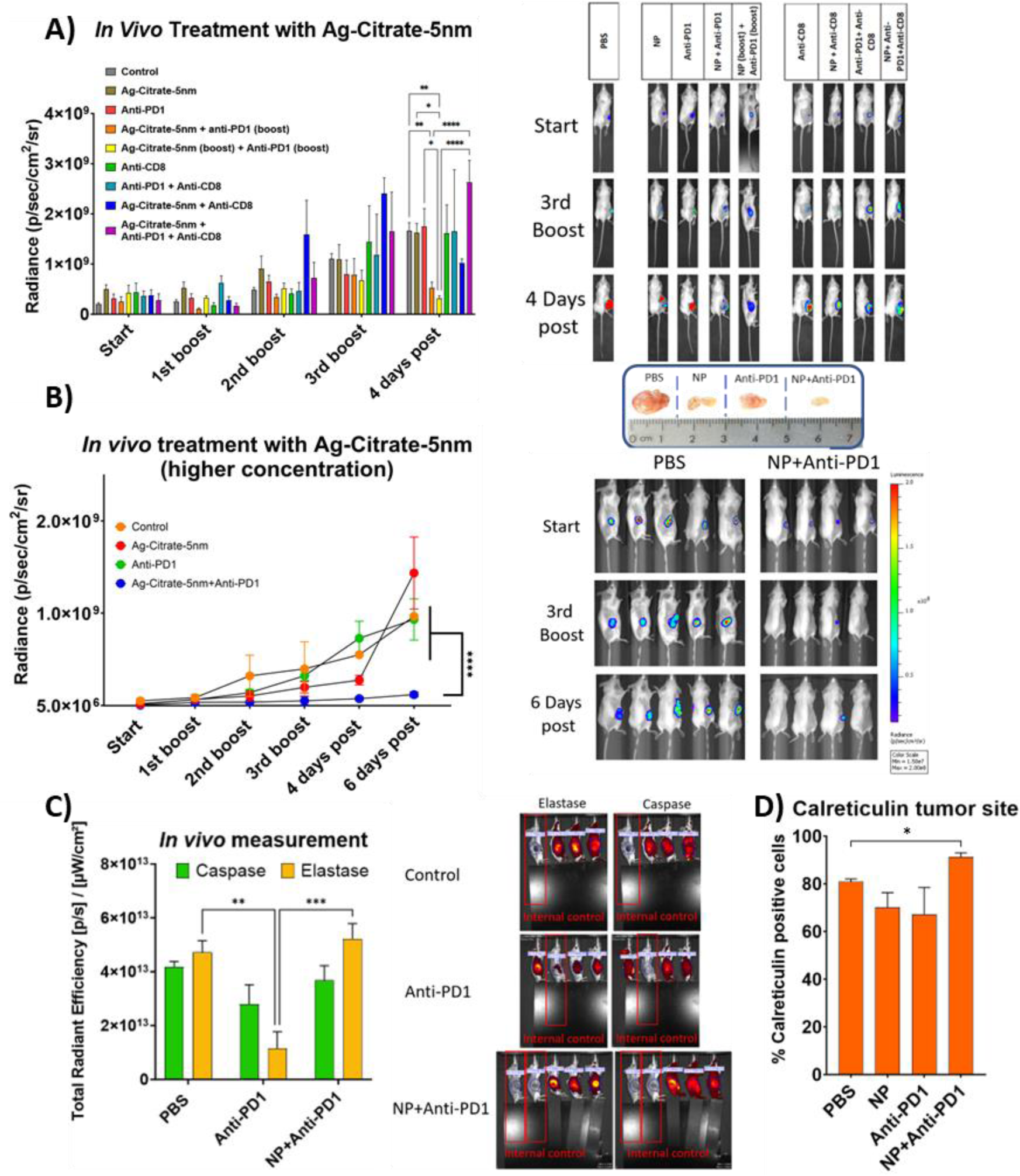
Therapeutic effects of Ag-citrate-5nm on Renca tumors and syngergy with anti-PD1 ICB. A) Relative photon flux (radiance) of Renca-luc+ tumors either injected a) peritumorally with PBS (100 μl/animal; control), b) peritumorally with Ag citrate-5 nm (20 μg/mouse), c) intraperitoneally with an anti-PD-1 antibody (150 μg/animal). The mice in the combination groups were injected peritumorally with Ag citrate-5 nm (20 μg/mouse) and intraperitoneally with anti-PD-1 antibody (150 μg/animal). The combination groups and the IT group were in total injected intraperitoneally with anti-PD-1 three times (boost) with an interval of 4 days between injections. Further control groups include 1) intraperitoneal administration of anti-CD8 (150 µg/ml), 2) anti-PD1 + anti-CD8, 3) Ag citrate-5 nm + anti-CD8 or Ag citrate-5 nm + anti-PD1 + anti-CD8. Representative bioluminescence images of mice with Renca Luc^+^ xenograft tumors in different treatment groups. These groups also include treatment with anti-CD8 antibody either as monotherapy, or in combination with NPs, anti-PD1 or both where anti-CD8 antibody (150 μg/animal) was administered intraperitoneally three times with an interval of 4 days between injections. B) Tumor flux of Renca Luc^+^ cells in photon per flux. Subcutaneous Renca tumors were treated with higher concentration of Ag-citrate-5nm (50µg/mouse) and Anti-PD1 (200µg/mouse). Representative bioluminescence images of mice with Renca Luc^+^ xenograft tumors in different treatment groups. Representative image of the tumors at the final timepoint. C) Bar graphs displaying the level of caspase and elastase at the tumor as measured through non-invasive optical imaging. Representative fluorescence images of mice with Renca Luc^+^ xenograft tumors in different treatment groups. D) Bar graphs showing the cancer cell-selective calreticulin translocation to the cell surface as evaluated through ImageStream based flow cytometry upon treatment with the various conditions. The results are presented as the mean of the animals/group ± SEM. The level of significance was indicated when appropriate (*:p< 0.05; **:p< 0.01; ***:p < 0.001).

To evaluate the role of cytotoxic (CD8^+^) T cells in the therapeutic potency of the Ag NPs in their combination therapy with ICB, experiments were repeated in the presence of anti-CD8 antibodies to deplete CD8^+^ T cells (**Figure 5A)**. The data reveal that depletion of CD8^+^ T cells largely impeded the therapeutic efficacy of the Ag-citrate-5nm and anti-PD1 combination group. However, despite the presence of CD8^+^ T cell neutralizing antibodies, a reduction in tumor growth was still apparent. This may be due to the direct cytotoxicity of the Ag NPs, although this is unlikely given that the NP only group did not reveal the same trend. A more likely explanation is that anti-PD1 therapy not only affected CD8^+^ T cells. Recent data have revealed that while PD1 is indeed not limited to T cells, but is also expressed in TAMs, making TAMs inactivate upon binding tumor cell-associated PD-L1. Treatment with an anti-PD-1 antibody therefore not only activates cytotoxic T cells, but was found to also affect TAMs so that they increase phagocytic potency against tumor cells(45). This study provides results which corroborate the findings of previous work done by Naatz et al. where they clearly showed therapeutic effects without relapse of tumor cells post-treatment with 6 % Fe-doped CuO NPs combined with myeloid-derived suppressor cell silencing(19). The combination group of Ag-citrate-5nm with anti-PD1 also showed a slight increase in elastase activity at the tumor site, indicating a more severe inflammatory process. Although the increased levels of neutrophil elastase correspond with increased inflammation, this is not favorable for tumor killing. Lerman *et al* showed that infiltrating myeloid cells exert protumorigenic effects via neutrophil elastase(46). Our results also showed that the combination group did not increase caspase activation in mice, indicating that the combination group did not induce apoptosis. These results are consistent with our findings that Ag citrate-5nm treatment in combination with anti-PD1 triggered significantly high calreticulin translocation at the tumor site and therefore suggest the induction of ICD, which is contrary to apoptosis as a non-immunogenic form of cell death.

To further investigate the effect of the combination group, we increased the concentration of Ag-citrate-5nm to 50µg/mouse (*n = 8*) and the anti-PD1 concentration to 200µg/mouse (*n = 8*). The results showed that at higher concentration of the therapeutic agents, tumor growth could be further suppressed (**Figure 5 B**).

### 3.6 Ag NPs and anti-PD1 ICB generate a potent anti-tumor immune response

To analyse the effect of different treatments in the activation of T cells, we stained tumor tissue sections with anti-CD8 and F4/80 antibodies to investigate which treatments increased the presence of cytotoxic T cells and TAMs at the tumor site, respectively. The control group showed an accumulation of cytotoxic T cells and macrophages at the tumor periphery, while treatment with AgNPs and anti-PD1 showed more infiltrated immune cells. The highest infiltration was clearly observed in the combination group (**Figure 6**).

**Figure 6.**
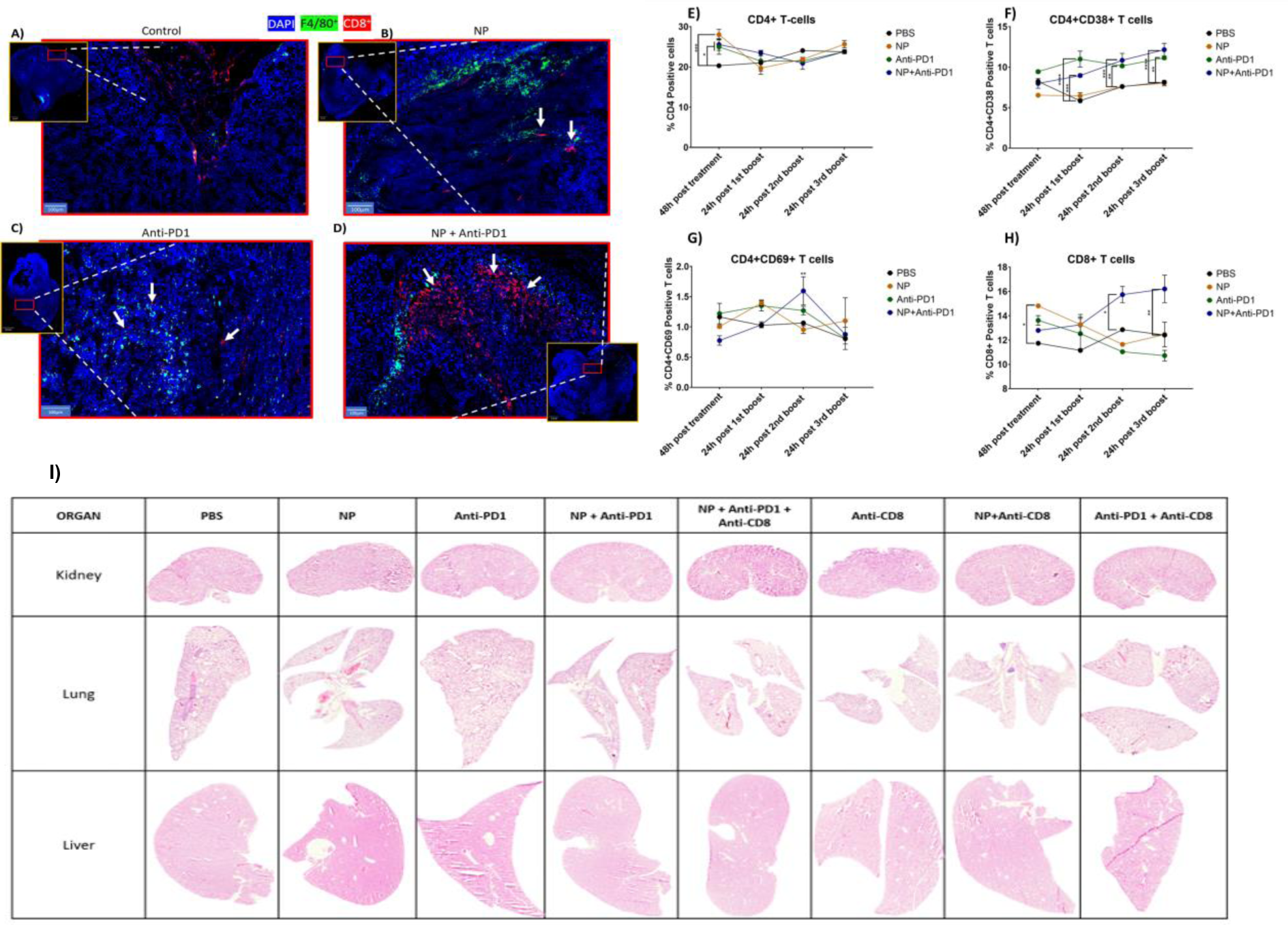
Effect of Ag-citrate-5nm, with or without IT, on immune cell activation in the spleen and tumor tissue. Representative fluorescence images of tumor tissue section obtained from Renca luc+ tumors treated with A) PBS, B) Ag-citrate-5nm monotherapy, C) anti-PD1 monotherapy or D) combination therapy. The images reveal tissue sections stained for F4/80 (green, TAMs), CD8 (CD8^+^ T cells, red) and counterstained with DAPI (cell nuclei, blue). Scale bars of 100 µm are indicated in the bottom left corner. Bar graphs displaying the levels of E) CD4+ T cells in the spleen, F) CD4^+^ CD38+ active T cells in the spleen, G) CD4^+^ CD69+ active T cells in the spleen or H) CD8+ T cells in the spleen. The results are presented as the normalized mean ± SEM in percentages related to the control group (PBS =100%). The level of significance was indicated when appropriate (*:p<0.05; **:p< 0.01; ***:p<0.001; ****:p<0.0001). I) Representative H&E stained images of kidney (top row), lung (middle row) and liver (bottom row) tissue sections of tumor-bearing mice treated with the respective agents indicated at the top.

To evaluate whether the therapies would be able to induce systemic immune responses, splenocytes were isolated from mice having received the different treatments. Flow cytometry analysis revealed significant increased levels of CD4+ T cells 48h after treatment with either NP, Anti-PD1 or combination of both compared to the control group. (**Figure 6E**). The combination group of NPs + Anti-PD1, resulting in an increase in CD8^+^ T cells present compared to the other groups (**Figure 6H)**. Splenocyte staining of the combination group also showed significant higher CD69^+^ T (p < 0.01) and CD38^+^ T-cell compared to control group (p < 0.0001) (**Figure 6F and 6G**). The increase in CD69^+^ T cells combined with an increase in CD8^+^ T cells only in the combination group, suggests that the combined therapy was able to generate anti-tumor immunity in the spleen, which would indicate the generation of a systemic anti-tumor effect. As CD69 is typically seen as an early marker of T cell activation(47), rather than CD38, which is a general marker for active cells, the shift towards more CD69^+^ T cells indicates the recent activation of T cells. This will likely affect both CD4^+^ and CD8^+^ T cells, but the increase in CD8^+^ T cells is in line with the local infiltration of CD8^+^ T cells in the tumor.

Together, these data reveal that only the combination group of Ag-citrate-5nm with anti-PD1 had more CD8^+^ in splenocyte staining. Splenocyte staining also showed a significantly high presence of CD69^+^ T cells, suggesting that the treatment activated the immune system. We could see that when the tumor was treated with Ag-citrate-5nm, there was higher level of immune cells present at the tumor site implying that Ag-citrate-5nm is capable of inducing cell death which attracts immune cells to the tumor site. These data support the results obtained for tumor growth and TME immunomodulation. Tumors treated with Ag-citrate-5nm and anti-PD1, showed significant lower tumor flux compared to the control group at day 30. Of interest, hematoxylin and eosin staining of vital organs revealed no structural damage to any of the organs (liver, lungs, kidney) studied (**Figure 6I**).

## 4 Conclusions

In the present study, we observed that among different metallic NPs and differently sized and coated AgNPs, Ag-citrate-5nm showed the highest cytotoxic effect. This was particularly true in Renca cells which were highly sensitive to Ag-citrate-5nm, where it induced ROS mediated cytotoxicity associated with a reduction in mitochondrial area and shrinkage of Renca cells. The cell death and cell stress induced by Ag-citrate-5nm resulted in translocation of calreticulin from the endoplasmic reticulum to the cell surface, which is an important marker of immunogenicity, and significantly increased the expression of several cytokines with potent immune modulating properties. Combined with classical anti-PD1 ICB, Ag-citrate-5nm resulted in highly significant tumor growth reduction, higher levels of tumor cell death, increased infiltration of immune cells at the tumor site, elevated calreticulin translocation, and neutrophil elastase activity, increased levels of newly activated CD4+ T cells and increased population of CD8+ T cells in the spleen. Overall, our study provides new insights into the use of Ag citrate-5nm NPs as a potential combination therapy with IT. Finally, we suggest that different concentrations, routes of administration, and regimens of combination therapy could be explored in the future to obtain the most effective treatment strategy for subcutaneous Renca tumors and to validate this strategy in orthotopic tumor models.

## Acknowledgements

This work was supported by the KU Leuven C3/20/090, C3/21/035, C24/18/101, and IDN/21/013 fundings acquired by Dr. Bella B. Manshian and the European research council (ERC) ERC StG 750973 funding acquired by Prof. Stefaan J. Soenen.

## Declaration of Interest statement

The authors declare that they have no known competing financial interests or personal relationships that could have appeared to influence the work reported in this paper.

## Notes

### Competing Interest Statement

The authors have declared no competing interest.

